## Supplementary Notes for "SparsePro: an efficient fine-mapping method integrating summary statistics and functional annotations"

### SparsePro Supplementary Notes

[3]McGill Genome Centre, Montreal, Canada

#### 1 Equivalence between the SuSiE IBSS algorithm and a paired mean field variational inference algorithm

Starting with the motivation to quantify uncertainty in selecting variants for constructing credible sets, SuSiE introduced a novel model that decomposes multivariate regression into a sum of univariate regressions [1]. This innovative approach led to the development of the IBSS algorithm, which enables efficient and accurate fine-mapping [1]. The computational efficiency of the IBSS algorithm can be better understood through the lens of dimensionality reduction. Specifically, introducing a sparse projection ($\mathbf{S}_{G\times K}$) on the high dimensional genotype matrix ($\mathbf{X}_{N\times G}$) can group correlated variants into $K$ effect groups. With the number of effect groups being much smaller than the number of total variants ($K << G$), inferring effect group-level causal configurations is less computationally expensive than exhaustively searching for variant-level causal configurations, leading to significantly improved efficiency.

Building upon this perspective, we provide an alternative sparse projection formulation of the SuSiE model and establish a connection between the IBSS algorithm and a well-studied paired mean field variational inference algorithm [2]. Here, we demonstrate their equivalence, with the aim of improving the understanding of both algorithms.

Specifically, without functional annotations, in the sparse projection formulation, we denote the sparse projection on genotype matrix as $\mathbf{S} = [\mathbf{s}_1, ..., \mathbf{s}_K]$ and the effect size vector as $\boldsymbol{\beta} = [\beta_1, ..., \beta_K]$ where $\mathbf{s}_k \sim \textbf{Multinomial}(1, \tilde{\boldsymbol{\pi}})$ is the sparse indicator for the variant compositions and $\beta_k \sim \mathcal{N}(0, \tau_\beta^{-1})$ is the corresponding effect size of the $k^{th}$ effect group. Under a linear model, for a continuous trait $\mathbf{y}$, we have $\mathbf{y} \sim \mathcal{N}(\mathbf{XS}\boldsymbol{\beta}, \tau_y^{-1}\mathbf{I})$.

Inference of the exact posterior distribution of the sparse projection is challenging. To address this, Titsias et al [2] proposed a paired mean field factorized variational family $q(\mathbf{S}, \boldsymbol{\beta}) = \prod_k q(\mathbf{s}_k, \beta_k) = \prod_k q(\mathbf{s}_k) q(\beta_k|\mathbf{s}_k)$ to approximate the posterior distribution. The proposed variational distribution maintains the dependency between $\mathbf{s}_k$ and $\beta_k$, and has been shown to closely resemble the mode and shape of the desired posterior distribution. This results in accurate estimations with significantly improved computational efficiency [2]. In the SuSiE IBSS algorithm, a similar approximation that maintains this dependency is achieved under the single-effect regression [1, 3].

Obtaining the optimal approximation under the variational inference framework involves minimizing the Kullback-Leibler (KL) divergence between the posterior distribution and the proposed variational distribution, which is equivalent to maximizing the evidence lower bound (ELBO), a lower bound on the log-likelihood of the data. We can achieve this by satisfying the conditions $\log q(\mathbf{s}_k, \beta_k) = E_{q(\mathbf{s}_{\setminus k}, \boldsymbol{\beta}_{\setminus k})}[\log p(\mathbf{y}, \mathbf{S}, \boldsymbol{\beta}|\mathbf{X})]$ where $E_{q(\mathbf{s}_{\setminus k}, \boldsymbol{\beta}_{\setminus k})}$ denotes the expectation with respect to the variational distribution excluding the $k^{th}$ component [4]. Next, we show that these conditions are equivalent to posterior inference under the single-effect regression in SuSiE [1, 3].

Based on the sparse projection formulation, we have:

$$\begin{aligned}\log p(\mathbf{y}, \mathbf{S}, \boldsymbol{\beta}|\mathbf{X}) &= \log p(\mathbf{y}|\mathbf{X}, \mathbf{S}, \boldsymbol{\beta}) + \sum_k \log p(\beta_k|\tau_\beta) + \sum_k \log p(\mathbf{s}_k|\tilde{\boldsymbol{\pi}}) \\ &= \frac{N}{2}\log\frac{\tau_y}{2\pi} - \frac{\tau_y}{2}(\mathbf{y} - \mathbf{X}(\sum_k \mathbf{s}_k\beta_k))^\top(\mathbf{y} - \mathbf{X}(\sum_k \mathbf{s}_k\beta_k)) \\ &\quad + \sum_k(\frac{1}{2}\log\frac{\tau_\beta}{2\pi} - \frac{\tau_\beta}{2}\beta_k^2) + \sum_k\sum_g s_{kg}\log\tilde{\pi}_g \end{aligned} \tag{1}$$

Denoting $\tilde{\boldsymbol{\beta}}_{\backslash k} = E_{q(\mathbf{s}_{\backslash k}\boldsymbol{\beta}_{\backslash k})}[\sum_{k\prime \neq k} \mathbf{s}_k \beta_k]$, the required conditions can be simplified as:

$$\log q(s_{kg} = 1, \mathbf{s}_{k\backslash g} = \mathbf{0}, \beta_k) = const - \frac{\tau_\beta}{2}\beta_k^2 - \frac{\tau_y}{2}\mathbf{X}_g^\top \mathbf{X}_g \beta_k^2 + \tau_y \beta_k \mathbf{X}_g^\top (\mathbf{y} - \mathbf{X}\tilde{\boldsymbol{\beta}}_{\backslash k}) + \log \tilde{\pi}_g \quad (2)$$

From which we have:

$$\begin{aligned}
& q(\beta_k | s_{kg} = 1, \mathbf{s}_{k\backslash g} = \mathbf{0}) \sim \mathcal{N}(\mu_{kg}^*, \tau_{kg}^*) \\
& \tau_{kg}^* = \tau_y \mathbf{X}_g^\top \mathbf{X}_g + \tau_\beta \\
& \mu_{kg}^* = \frac{\tau_y}{\tau_{kg}^*}\mathbf{X}_g^\top (\mathbf{y} - \mathbf{X}\tilde{\boldsymbol{\beta}}_{\backslash k})
\end{aligned}$$

By integrating out $\beta_k$ in Equation (2), we have:

$$\log q(s_{kg} = 1, \mathbf{s}_{k\backslash g} = \mathbf{0}) = \log \tilde{\pi}_g - \frac{1}{2}\log \frac{\tau_{kg}^*}{2\pi} + \frac{1}{2}\tau_{kg}^* \mu_{kg}^{*2} + const$$

Denoting the posterior probability for the $g^{th}$ variant being included in the $k^{th}$ effect group as $\gamma_{kg}^*$, we have:

$$\begin{aligned}
\gamma_{kg}^* := q(s_{kg} = 1, \mathbf{s}_{k\backslash g} = \mathbf{0}) &= \frac{\exp(\log \tilde{\pi}_g - \frac{1}{2}\log \tau_{kg}^* + \frac{1}{2}\tau_{kg}^* \mu_{kg}^{*2}))}{\sum_{g\prime} \exp(\log \tilde{\pi}_{g\prime} - \frac{1}{2}\log \tau_{kg\prime}^* + \frac{1}{2}\tau_{kg}^* \mu_{kg\prime}^{*2}))} \\
&= \frac{\tilde{\pi}_g \sqrt{\exp(\tau_{kg}^* \mu_{kg}^{*2})\tau_{kg}^{*-1}}}{\sum_{g\prime} \tilde{\pi}_{g\prime} \sqrt{\exp(\tau_{kg\prime}^* \mu_{kg\prime}^{*2})\tau_{kg\prime}^{*-1}}}
\end{aligned}$$

It is important to note that the paired mean field variational approximation of the posterior distributions for effect sizes are the same as calculated under the single-effect regression in SuSiE [1]. The posterior probabilities for the sparse projection are analogous to prior weighted Bayes Factors in SuSiE, without additional normalizing factors [1].

### 2 Hyperparameter estimation

There are two important prior hyperparameters that impact fine-mapping results: $\tau_\beta$ for effect sizes and $\tau_y$ for residual variance. SuSiE employs an empirical Bayes approach to iteratively infer causal variants and estimate these hyperparamters [1]. In this approach, hyperparameter estimation depends on inference of causal variants, which can lead to local optima, especially when the parameter space is large. To mitigate this issue, we propose to estimate these parameters outside of the fine-mapping algorithm. Interestingly, both hyperparameters are closely related to local heritability, which can be estimated without the knowledge of causal variants. Specifically, we can obtain the local heritability ($\hat{h}^2$) in a locus as well as per-variant heritability ($\hat{h}_v^2$) with the HESS [5] estimator using GWAS summary statistics, and set the hyperparameters with: $\tau_\beta^{-1} = \hat{h}_v^2$ and $\tau_y^{-1} = 1 - \hat{h}^2$. Shi et al [5] showcased that the HESS estimator accurately and robustly estimates local heritabilities for a variety of genetic architectures, making it suitable for hyperparameter estimation.

In simulations, we have observed that incorporating local heritability-based hyperparameter estimation can improve power for fine-mapping (**Supplementary Table** S2) while maintaining calibration of PIP (**Supplementary Figure** S8). By applying this strategy to SuSiE, we have observed substantial improvements in performance (**Supplementary Table** S2). For example, in the simulation setting with K = 5 and W = 2, SuSiE+HESS outperformed the original SuSiE by identifying a greater number of true causal variants with higher PIP values (**Supplementary Figure** S11). Moreover, the variant-level PIP obtained from SuSiE+HESS and SparsePro- were highly similar (**Supplementary Figure** S12), both achieving an overall AUPRC of 0.91 (**Supplementary Table** S2).

### 3 Posterior summaries

Summarizing posterior probabilities is crucial for interpreting results from fine-mapping algorithms. In SuSiE, for each single-effect regression, a candidate $\rho$-level credible set is constructed to summarize its posterior probabilities [1]. Specifically, variants are added in the $\rho$-level candidate credible set in descending order of their posterior probabilities [1, 6]. However, if the statistical support is weak, uninformative variants with small posterior probabilities may also be included into the set to meet the nominal coverage threshold. To address this issue, SuSiE uses a purity metric (minimum absolute correlation between pairs of variants within a set) to remove candidate credible sets that cannot attain the nominal coverage $\rho$ with informative variants [1]. However, purity is a complex metric, as it involves an interplay of $\rho$ and LD tightness. In practice, it might be challenging to find the appropriate threshold.

As an alternative, we propose to only summarize effect groups with attainable coverage greater than $\rho$ to $\rho$-level credible sets, avoiding the need of purity-based filtering. Specifically, we define the attainable coverage of the $k^{th}$ effect group as:

$$c_k = \sum_g \gamma_{kg}$$

with

$$\gamma_{kg} = \begin{cases} \gamma^*_{kg}, & \text{if } \gamma^*_{kg} = \max\left(\gamma^*_{1g}, \ldots, \gamma^*_{Kg}\right) \\ 0, & \text{otherwise} \end{cases}$$

This definition takes advantage of the fact that a variant can only contribute informatively to at most one effect group. Essentially, if a variant has been represented by one effect group with high posterior probability, it will be removed from consideration in other effect groups since its effect has already been accounted for. Consequently, the posterior probabilities for causal variants are always high in one effect group and negligible in other effect groups. However, if there are no actual causal signals, this approach may lead to one credible set including a large number of variants. To address this issue, we additionally use an entropy-based threshold to remove this uninformative credible set. Specifically, entropy measures the level of uncertainty in a probability distribution [7] and in the context of fine-mapping, entropy corresponds to the logarithm of the minimum number of variants required to represent each effect group. We set the default cutoff as $\log 20$, corresponding to a upper limit of 20 variants in tight LD with each other in a legitimate credible set. Users can adjust this value based on the level of LD tightness observed in their genotype data.

In simulations, this attainable coverage-based approach resulted in improved set-level summaries. Across different settings, the credible sets obtained from SuSiE+SparsePro (SuSiE with local heritability-based hyperparameter estimation combined with the posterior summaries proposed in SparsePro) exhibited a similar coverage with a higher power and smaller size compared to SuSiE+HESS (SuSiE with local heritability-based hyperparameter estimation) (**Supplementary Figure** S9).

Additionally, similar to SuSiE, we use PIP to summarize posterior probabilities at the variant-level. For the $g^{th}$ variant, we define the PIP value as:

$$PIP_g = 1 - \Pi_k(1 - \gamma_{kg}) = 1 - (1 - \max\left(\gamma_{1g}, \ldots, \gamma_{Kg}\right)) = \max\left(\gamma^*_{1g}, \ldots, \gamma^*_{Kg}\right)$$

In simulations, this maximization-based approach yields similar variant-level summaries as the multiplication-based approach used in SuSiE (**Supplementary Figure** S13), as they are equivalent when the negligible posterior probabilities are disregarded. However, in practice, the maximization-based approach is more computationally efficient than the multiplication-based approach.

### 4 Adaptation to summary statistics

The information in individual-level genotype $\mathbf{X}$ and phenotype data $\mathbf{y}$ are used in the form of $\mathbf{X}^\top\mathbf{X}$ and $\mathbf{X}^\top\mathbf{y}$ throughout the algorithm. These statistics can be derived from publicly available GWAS summary statistics z-scores (the ratio of per-variant effect size estimate to its standard error) and matched LD information (estimates of pairwise variant-variant Pearson correlation coefficient). With standardized genotypes and phenotypes, we can derive the required quantities using summary-level data $\mathbf{X}^\top\mathbf{X} = N * \mathbf{LD}$ and $\mathbf{X}^\top\mathbf{y} = \sqrt{N}\mathbf{z}$, where $N$ is the sample size, $\mathbf{LD}$ is the variant correlation matrix and $\mathbf{z}$ stands for z-scores in GWAS summary statistics. For binary traits, we can use z-scores derived from log-odds ratios and their standard errors to approximate the required quantities.

### 5 Functionally-informed prior

If relevant functional annotations are available, we can incorporate them in statistical fine-mapping to further prioritize causal variants via a functionally-informed prior:

$$\tilde{\boldsymbol{\pi}} = softmax(\mathbf{A}\mathbf{w})$$

where $\mathbf{A}_{G\times M}$ represents the annotation matrix and $\mathbf{w}_{M\times 1}$ is the vector of enrichment weight to be estimated. With annotation information incorporated into the log likelihood function (1), the objective function becomes:

$$\begin{aligned} ELBO &= const + \sum_k \sum_g \gamma^*_{kg} \log \tilde{\pi}_g \\ &= const + \sum_k \sum_g \gamma^*_{kg} \log \frac{\exp(\mathbf{A}_g \mathbf{w})}{\sum_g \exp(\mathbf{A}_g \mathbf{w})} \\ &= const + \sum_k \sum_g \gamma^*_{kg} [\mathbf{A}_g \mathbf{w} - \log(\sum_g \exp(\mathbf{A}_g \mathbf{w}))] \end{aligned}$$

To derive closed-form estimates for $\mathbf{w}$, we can take the derivatives of the objective function with respect to $\mathbf{w}$. For the $m^{th}$ element of $\mathbf{w}$, we have:

$$\begin{aligned} \frac{\partial ELBO}{\partial w_m} &= \sum_k \sum_g \gamma^*_{kg} [A_{gm} - \frac{\sum_g A_{gm} \exp(\mathbf{A}_g \mathbf{w}))}{\sum_g \exp(\mathbf{A}_g \mathbf{w}))}] \\ &= \sum_k \sum_g \gamma^*_{kg} [A_{gm} - \frac{\sum_g A_{gm} \exp(A_{gm} w_m) \exp(\sum_{m' \neq m} A_{gm'} w_{m'}))}{\sum_g \exp(A_{gm} w_m) \exp(\sum_{m' \neq m} A_{gm'} w_{m'}))}] \\ &= \sum_k \sum_g \gamma^*_{kg} [A_{gm} - \frac{\sum_g A_{gm} \exp(A_{gm} w_m) softmax(\sum_{m' \neq m} A_{gm'} w_{m'})}{\sum_g \exp(A_{gm} w_m) softmax(\sum_{m' \neq m} A_{gm'} w_{m'})}] \\ &= r_1 - (r_1 + r_0) \frac{k_1 e^{w_m}}{k_1 e^{w_m} + k_0} \end{aligned}$$

where

$$\begin{aligned} k_1 &= \sum_g [A_{gm} = 1] softmax(\sum_{m' \neq m} A_{gm'} w_{m'}) \\ k_0 &= \sum_g [A_{gm} = 0] softmax(\sum_{m' \neq m} A_{gm'} w_{m'}) \\ r_1 &= \sum_{k,g} [A_{gm} = 1] \gamma^*_{kg} \\ r_0 &= \sum_{k,g} [A_{gm} = 0] \gamma^*_{kg} \end{aligned}$$

Solving for $w_m$, we have:

$$w_m = \log \left( \frac{r_1 / r_0}{k_1 / k_0} \right)$$

Through iterating over all annotations, upon convergence, we obtain a joint estimate of the enrichment weight vector $\mathbf{w}$ that can be used to derive functionally-informed priors.

### 6 A G-test for selection of relevant annotations

Incorporating annotation information might not always be beneficial to fine-mapping. Therefore, it may be desirable to select relevant annotations. We have observed that the estimates for enrichment weights are particularly informative when considering a single binary annotation. Specifically, the enrichment weight for this annotation simplifies to

$$w = \log\left(\frac{r_1/r_0}{k_1/k_0}\right)$$

with

$$k_1 = \sum_g [A_{gm} = 1], \text{ the total number of variants with this annotation}$$
$$k_0 = \sum_g [A_{gm} = 0], \text{ the total number of variants without this annotation}$$
$$r_1 = \sum_{k,g} [A_{gm} = 1]\gamma^*_{kg}, \text{ the total number of causal variants with this annotation}$$
$$r_0 = \sum_{k,g} [A_{gm} = 0]\gamma^*_{kg}, \text{ the total number of causal variants without this annotation}$$

This enrichment weight is analogous to a relative risk estimate in a $2 \times 2$ contingency table. To calculate its standard error, we can leverage the standard error of a relative risk:

$$se(w) = \sqrt{\frac{1}{r_1} + \frac{1}{r_0} - \frac{1}{k_1} - \frac{1}{k_0}}$$

The statistical significance of functional enrichment can also be assessed with the log likelihood ratio test (G-test) [8]. By applying the G-test, we can identify suitable functional information to be used in deriving functionally-informed priors.
