## Supplementaru Figures for "SparsePro: an efficient fine-mapping method integrating summary statistics and functional annotations"

### Slide 1
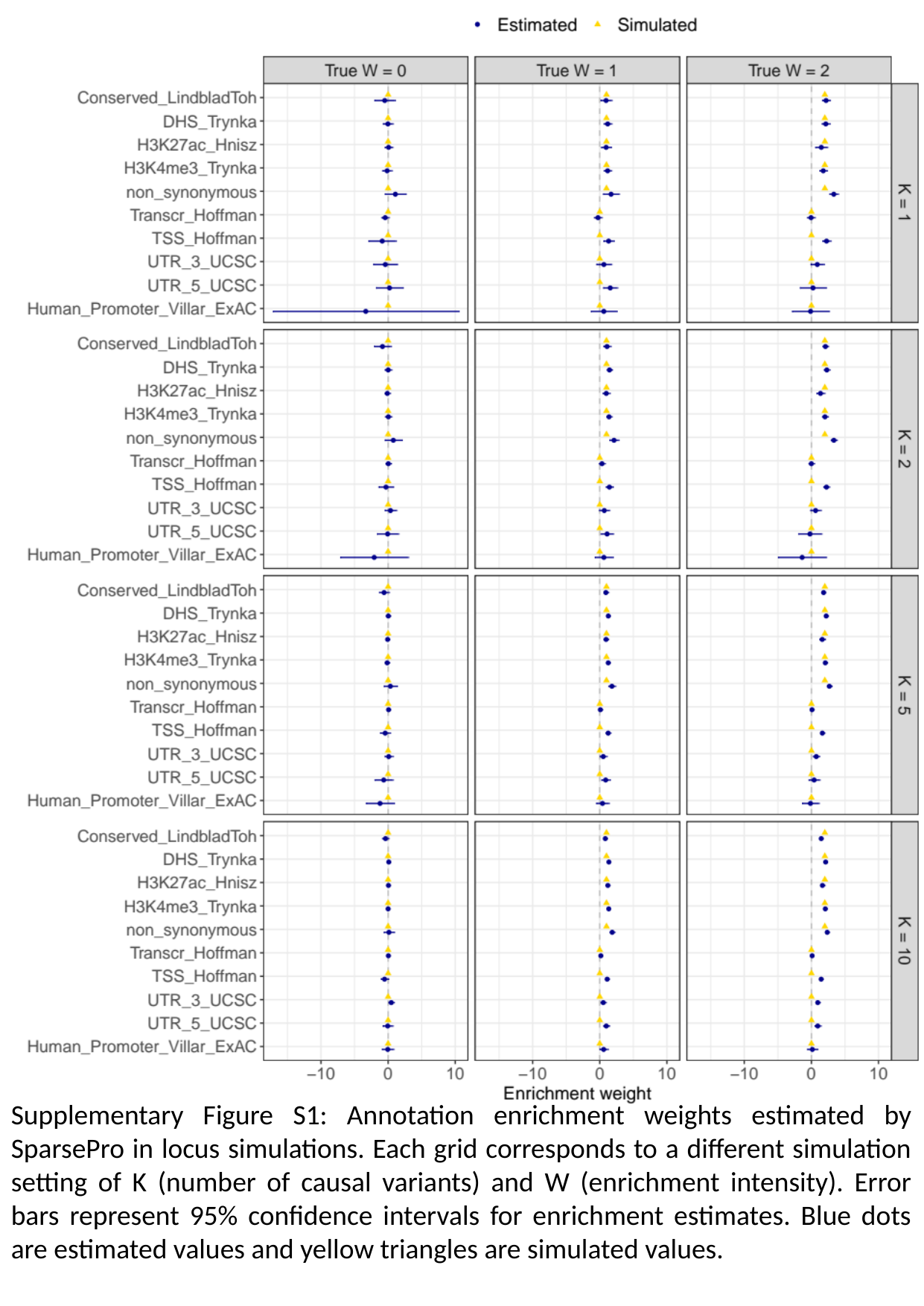

Supplementary Figure S1: Annotation enrichment weights estimated by SparsePro in locus simulations. Each grid corresponds to a different simulation setting of K (number of causal variants) and W (enrichment intensity). Error bars represent 95% confidence intervals for enrichment estimates. Blue dots are estimated values and yellow triangles are simulated values.

### Slide 2
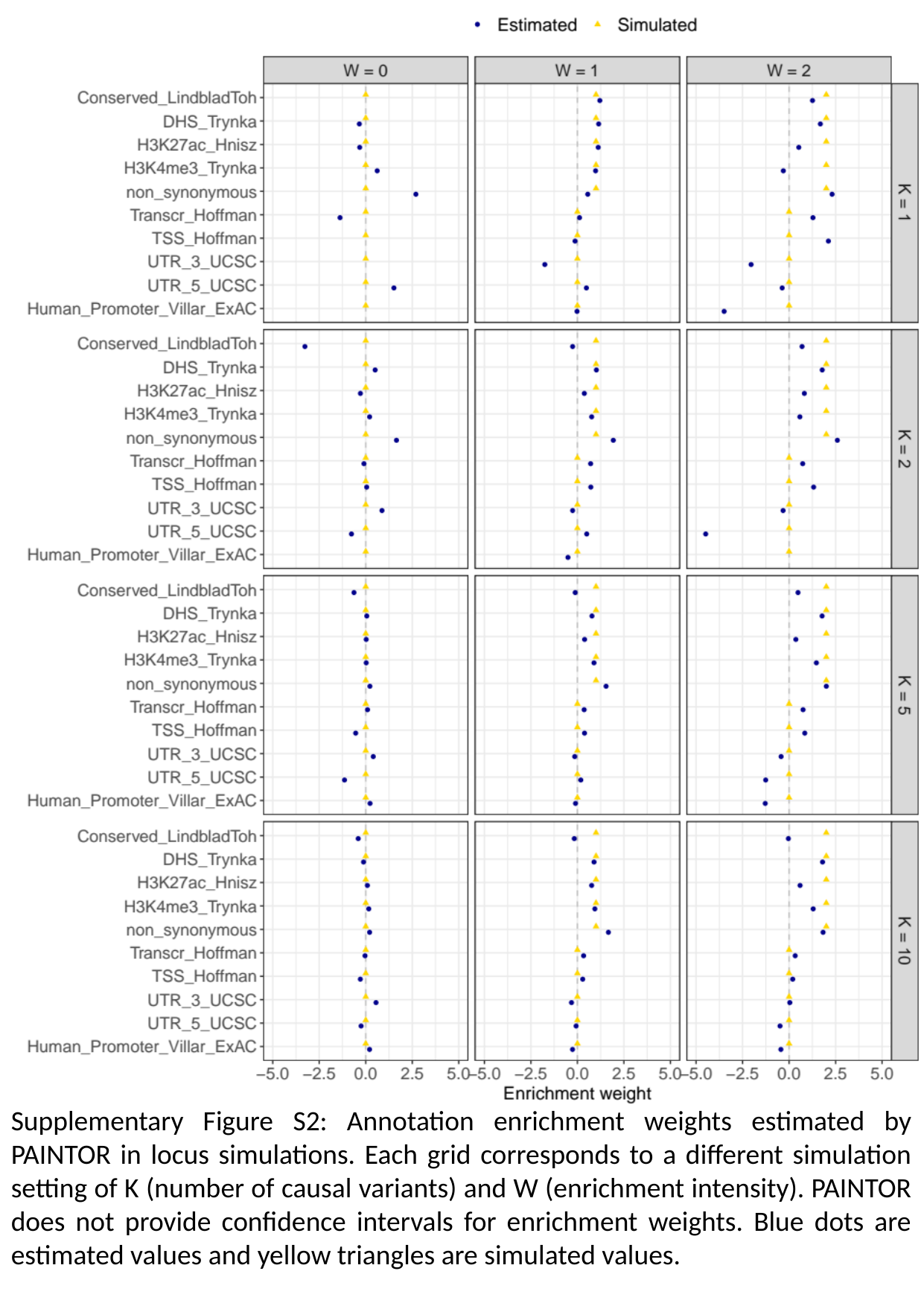

Supplementary Figure S2: Annotation enrichment weights estimated by PAINTOR in locus simulations. Each grid corresponds to a different simulation setting of K (number of causal variants) and W (enrichment intensity). PAINTOR does not provide confidence intervals for enrichment weights. Blue dots are estimated values and yellow triangles are simulated values.

### Slide 3
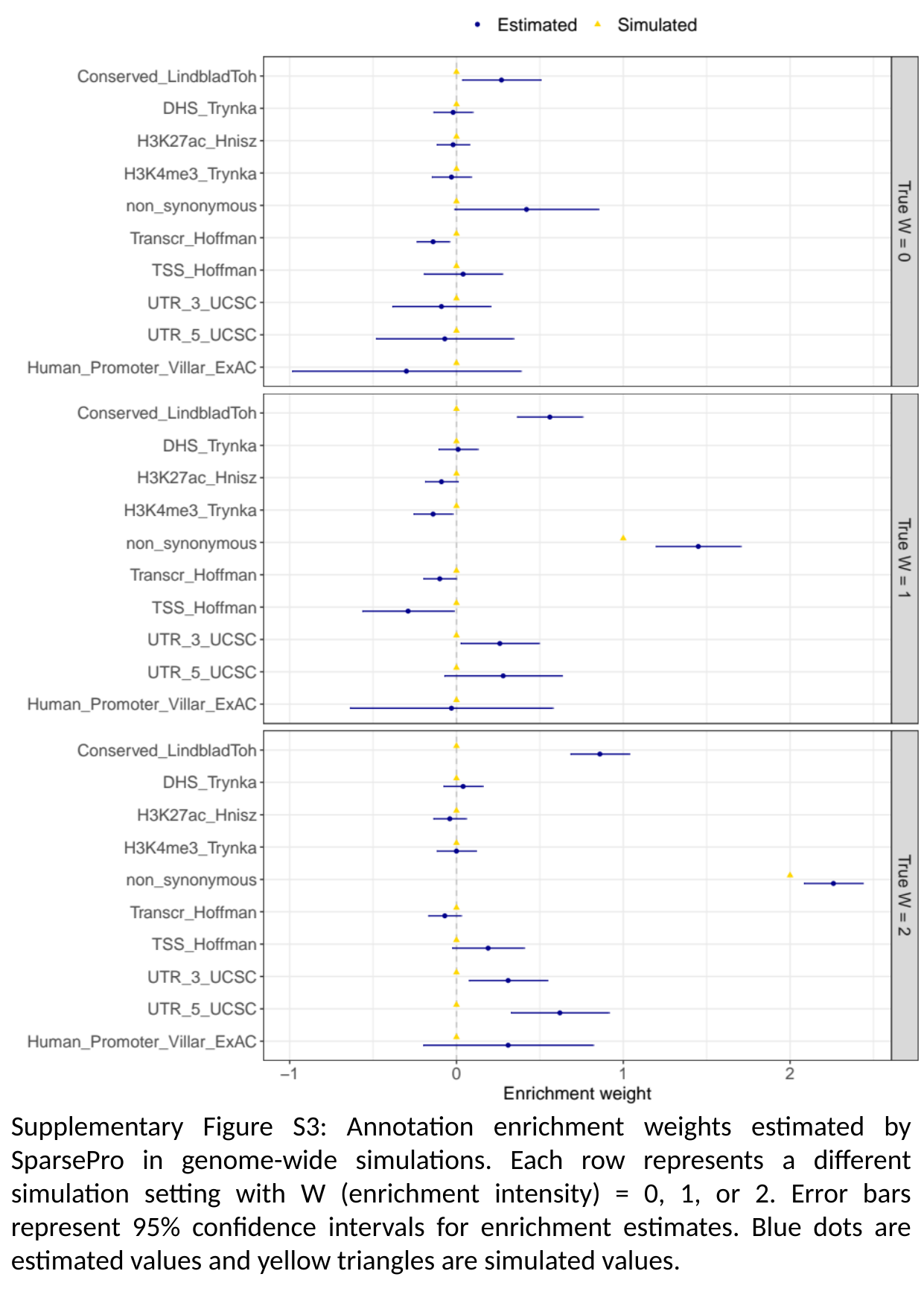

Supplementary Figure S3: Annotation enrichment weights estimated by SparsePro in genome-wide simulations. Each row represents a different simulation setting with W (enrichment intensity) = 0, 1, or 2. Error bars represent 95% confidence intervals for enrichment estimates. Blue dots are estimated values and yellow triangles are simulated values.

### Slide 4
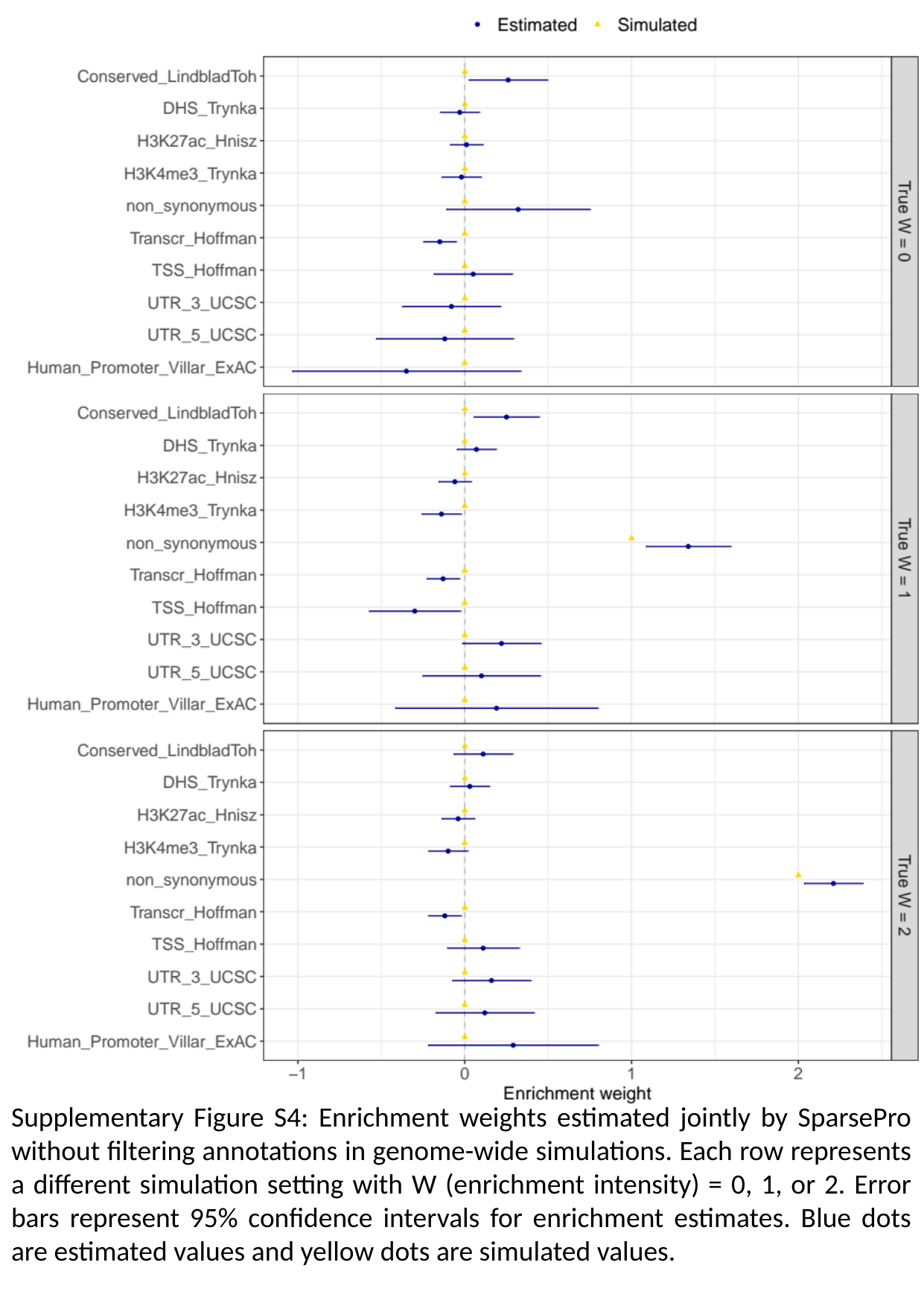

Supplementary Figure S4: Enrichment weights estimated jointly by SparsePro without filtering annotations in genome-wide simulations. Each row represents a different simulation setting with W (enrichment intensity) = 0, 1, or 2. Error bars represent 95% confidence intervals for enrichment estimates. Blue dots are estimated values and yellow dots are simulated values.

### Slide 5
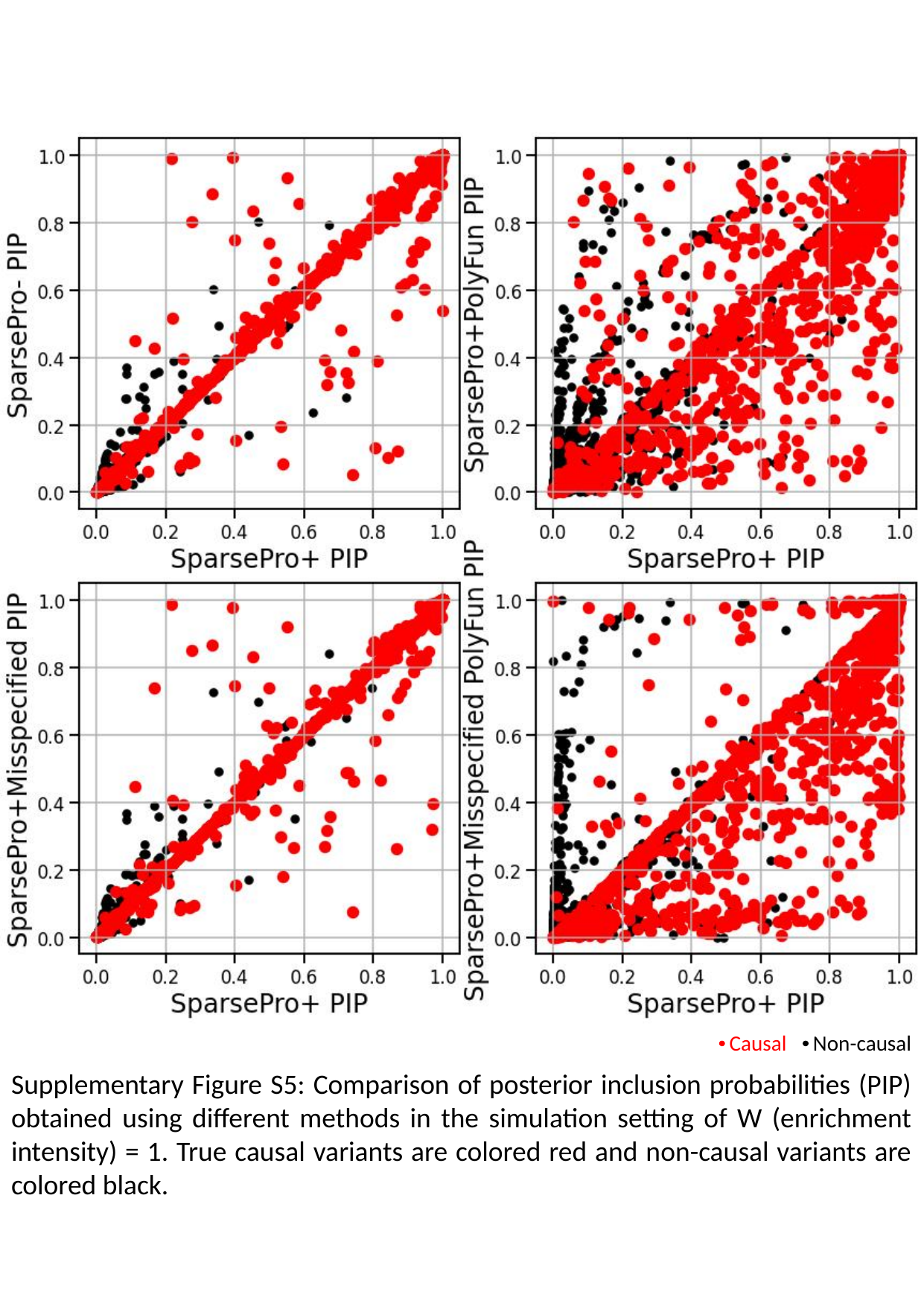

Causal
Non-causal
Supplementary Figure S5: Comparison of posterior inclusion probabilities (PIP) obtained using different methods in the simulation setting of W (enrichment intensity) = 1. True causal variants are colored red and non-causal variants are colored black.

### Slide 6
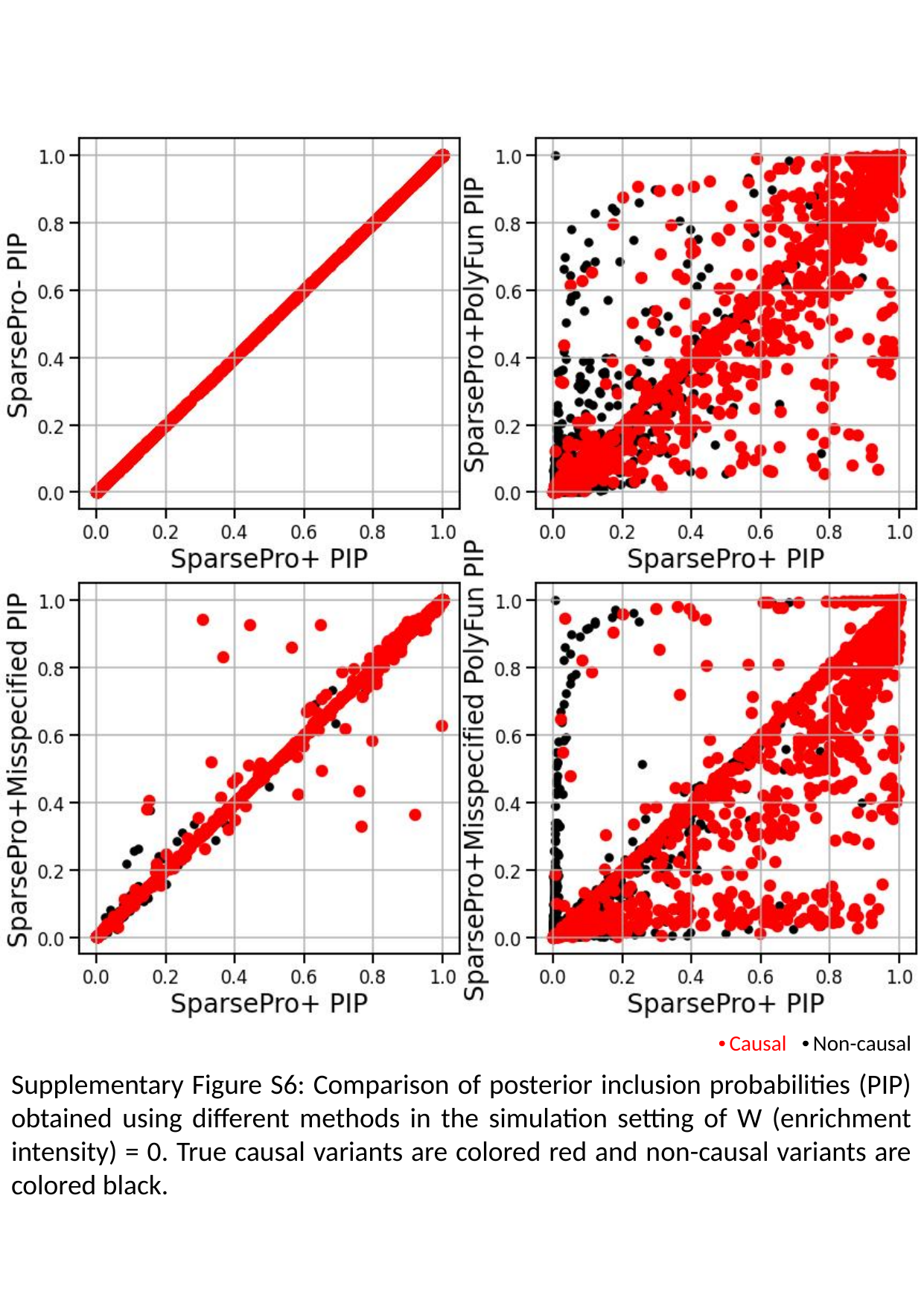

Causal
Non-causal
Supplementary Figure S6: Comparison of posterior inclusion probabilities (PIP) obtained using different methods in the simulation setting of W (enrichment intensity) = 0. True causal variants are colored red and non-causal variants are colored black.

### Slide 7
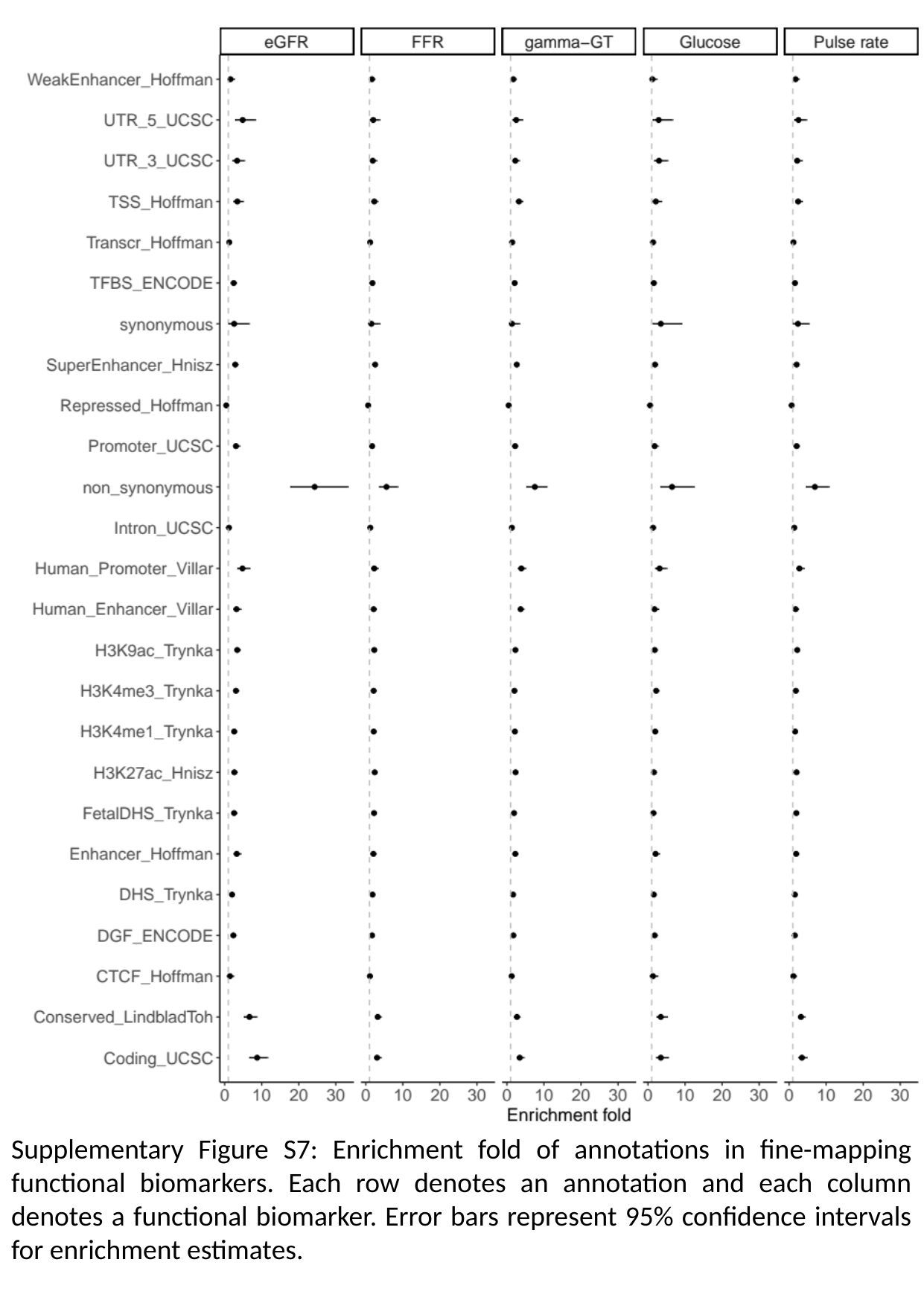

Supplementary Figure S7: Enrichment fold of annotations in fine-mapping functional biomarkers. Each row denotes an annotation and each column denotes a functional biomarker. Error bars represent 95% confidence intervals for enrichment estimates.

### Slide 8
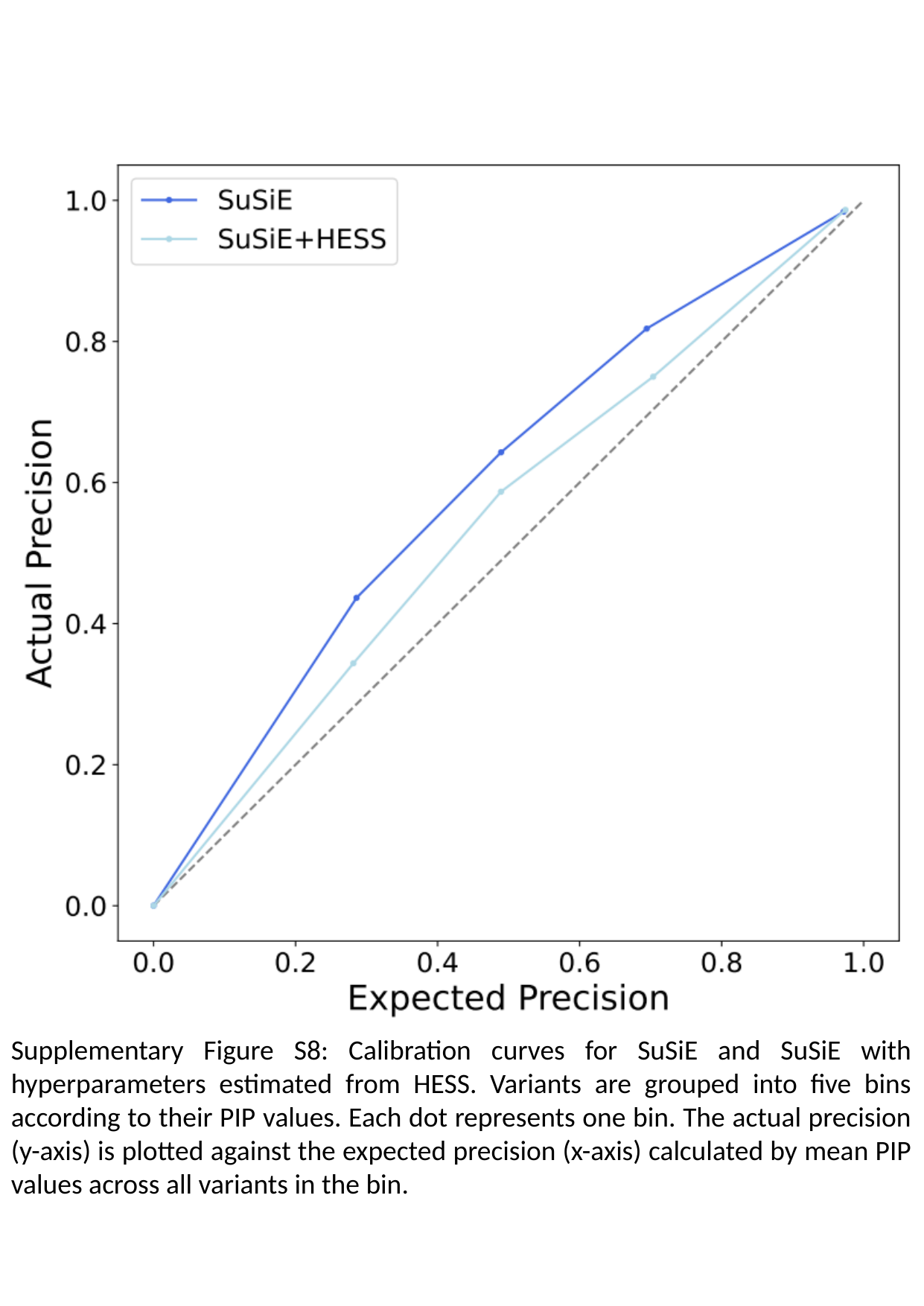

Supplementary Figure S8: Calibration curves for SuSiE and SuSiE with hyperparameters estimated from HESS. Variants are grouped into five bins according to their PIP values. Each dot represents one bin. The actual precision (y-axis) is plotted against the expected precision (x-axis) calculated by mean PIP values across all variants in the bin.

### Slide 9
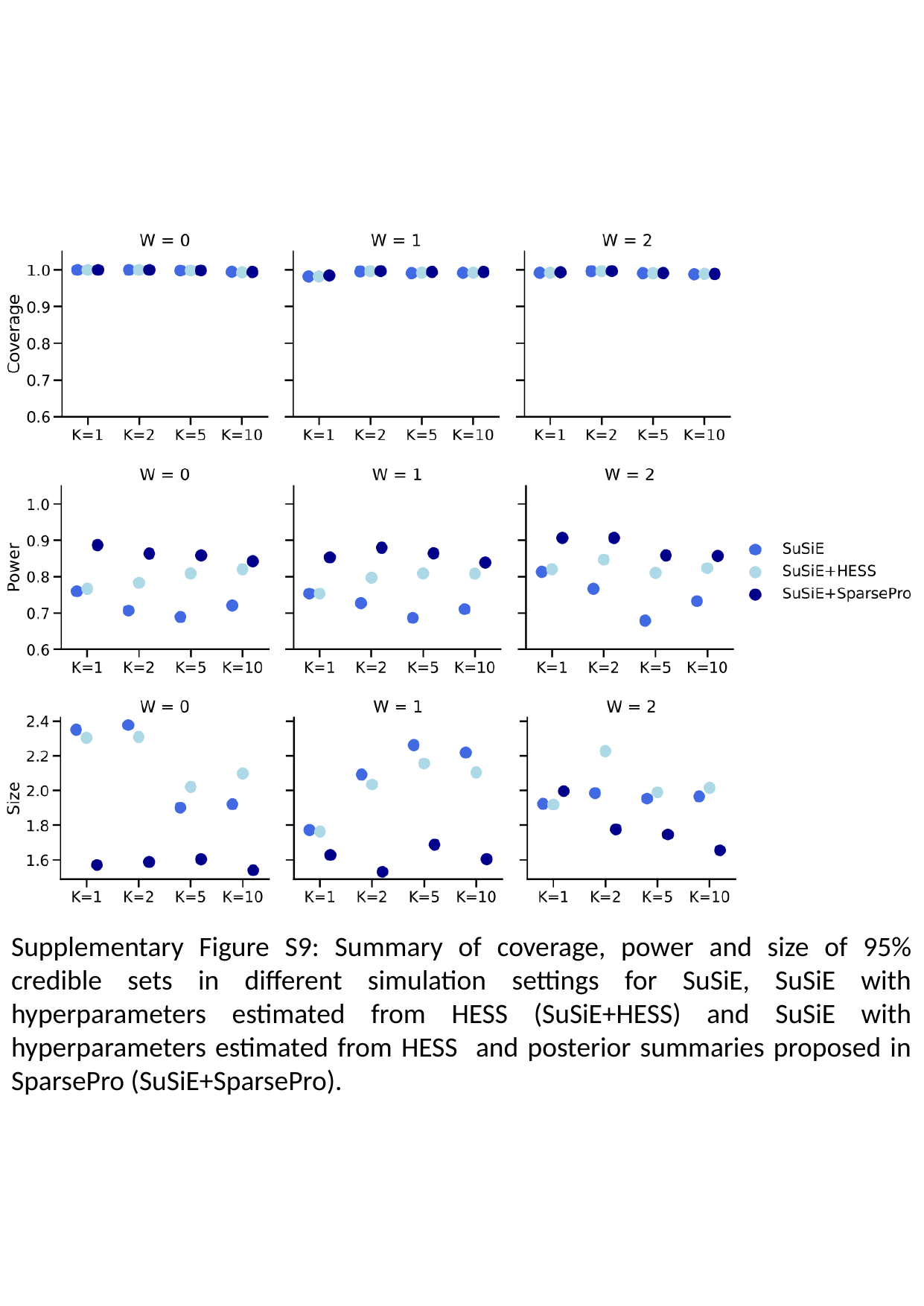

Supplementary Figure S9: Summary of coverage, power and size of 95% credible sets in different simulation settings for SuSiE, SuSiE with hyperparameters estimated from HESS (SuSiE+HESS) and SuSiE with hyperparameters estimated from HESS and posterior summaries proposed in SparsePro (SuSiE+SparsePro).

### Slide 10
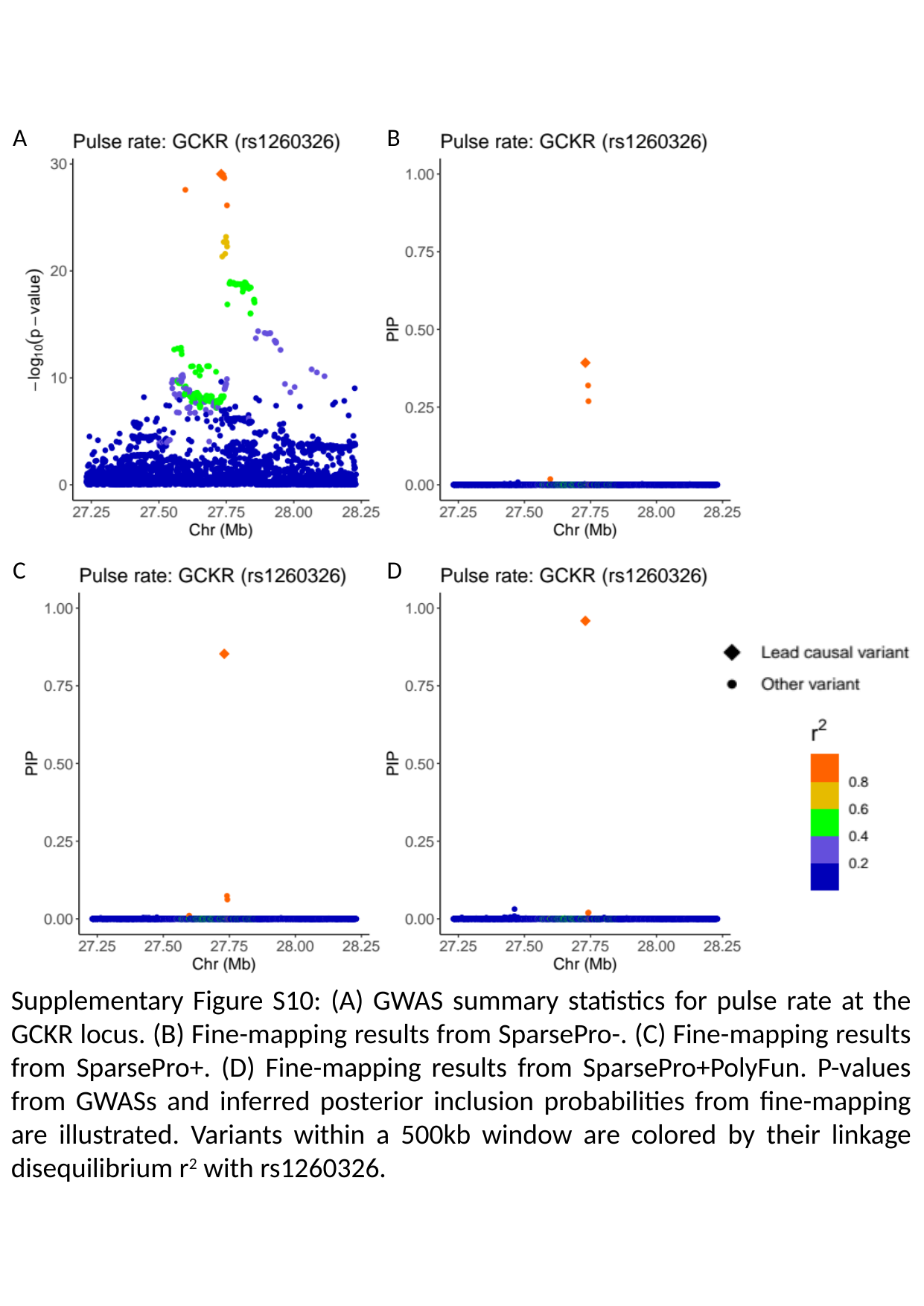

A
B
C
D

### Slide 11
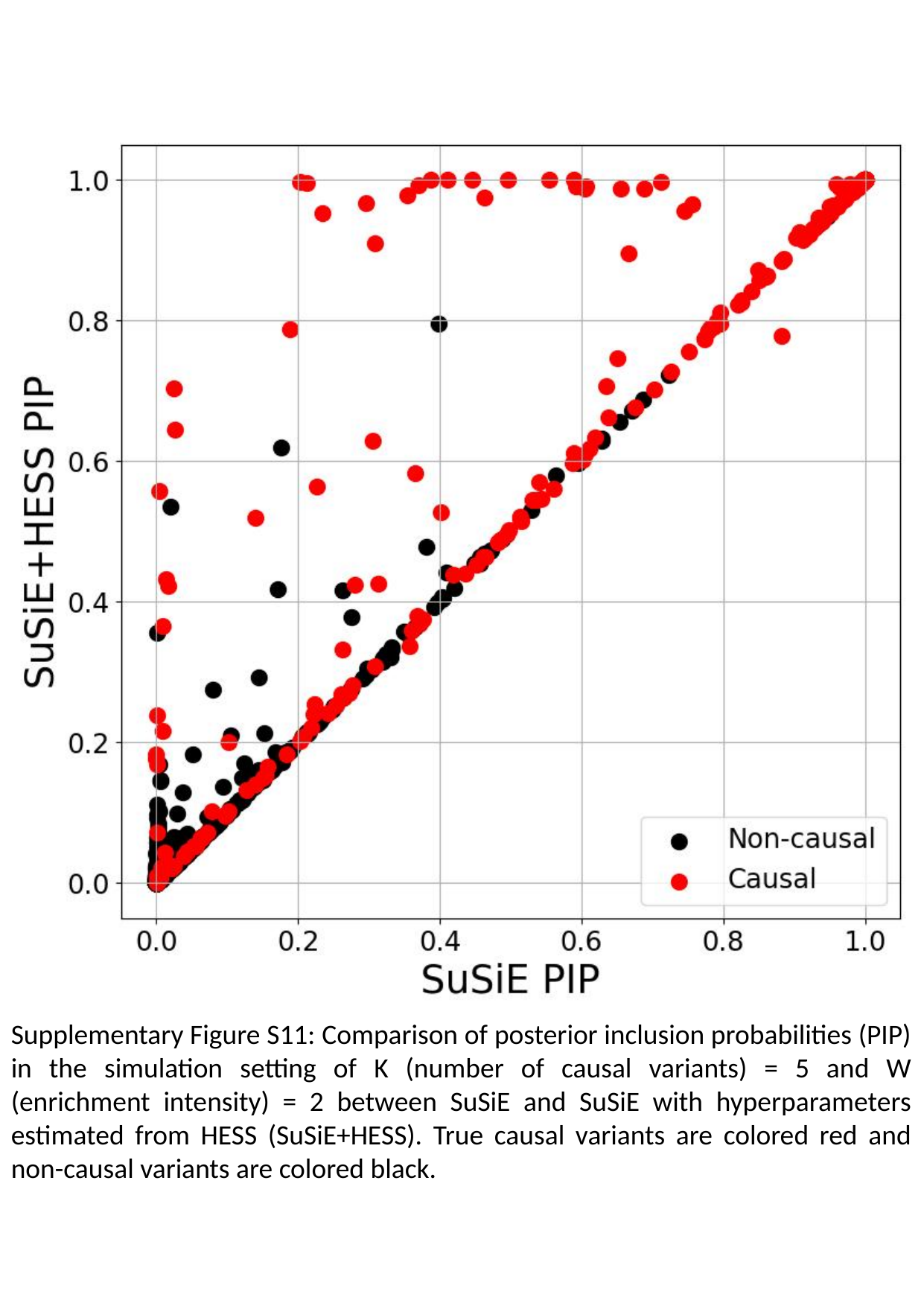

Supplementary Figure S11: Comparison of posterior inclusion probabilities (PIP) in the simulation setting of K (number of causal variants) = 5 and W (enrichment intensity) = 2 between SuSiE and SuSiE with hyperparameters estimated from HESS (SuSiE+HESS). True causal variants are colored red and non-causal variants are colored black.

### Slide 12
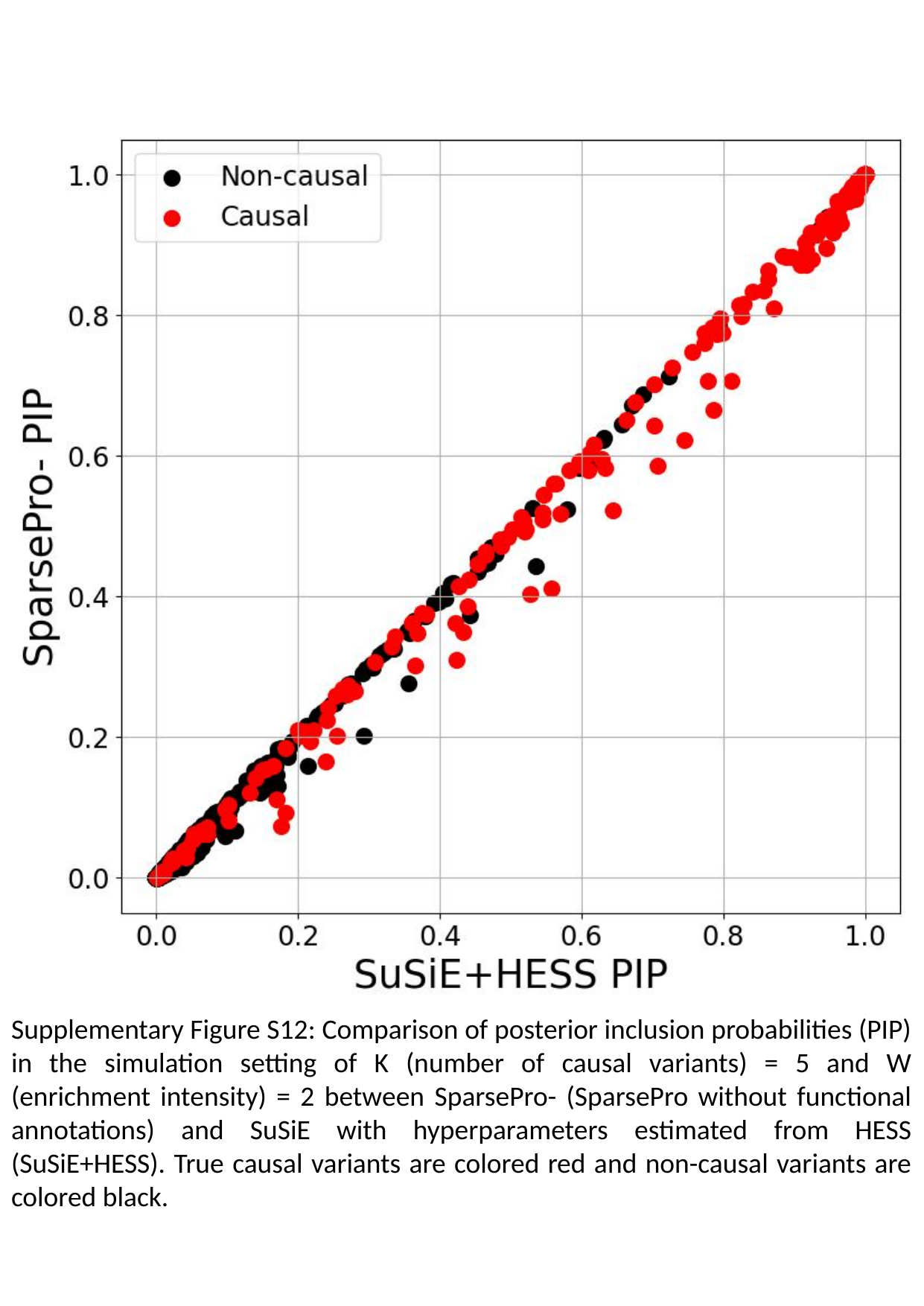

Supplementary Figure S12: Comparison of posterior inclusion probabilities (PIP) in the simulation setting of K (number of causal variants) = 5 and W (enrichment intensity) = 2 between SparsePro- (SparsePro without functional annotations) and SuSiE with hyperparameters estimated from HESS (SuSiE+HESS). True causal variants are colored red and non-causal variants are colored black.

### Slide 13
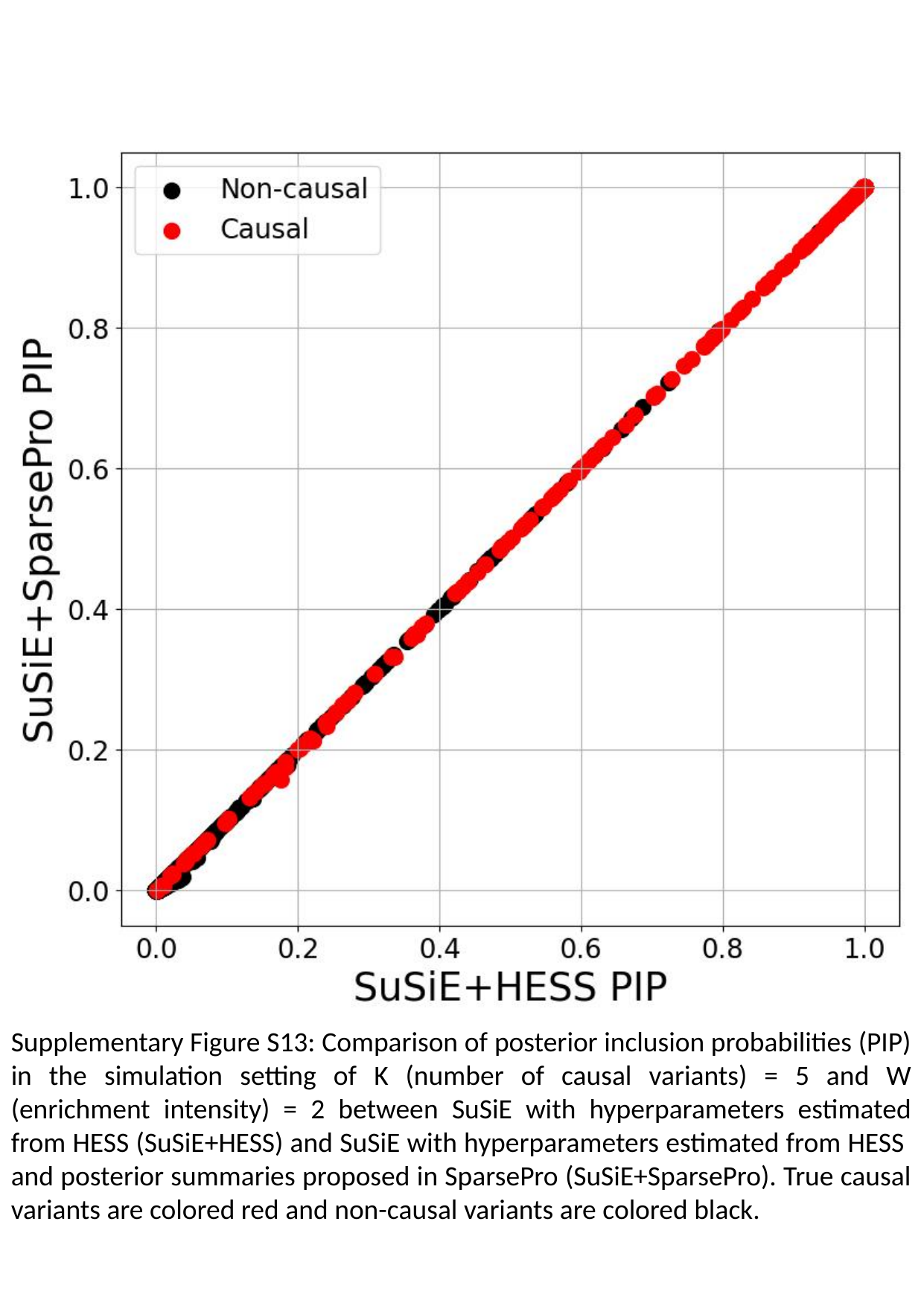

Supplementary Figure S13: Comparison of posterior inclusion probabilities (PIP) in the simulation setting of K (number of causal variants) = 5 and W (enrichment intensity) = 2 between SuSiE with hyperparameters estimated from HESS (SuSiE+HESS) and SuSiE with hyperparameters estimated from HESS and posterior summaries proposed in SparsePro (SuSiE+SparsePro). True causal variants are colored red and non-causal variants are colored black.
